# Genomic prediction informed by biological processes expands our understanding of the genetic architecture underlying free amino acid traits in dry *Arabidopsis* seeds

**DOI:** 10.1101/272047

**Authors:** Sarah D. Turner-Hissong, Kevin A. Bird, Alexander E. Lipka, Elizabeth G. King, Timothy M. Beissinger, Ruthie Angelovici

**Author notes:** Corresponding author: Ruthie Angelovici, 371D Christopher S. Bond Life Sciences, University of Missouri, Columbia, MO 65201.

## Abstract

Plant growth, development, and nutritional quality depends upon amino acid homeostasis, especially in seeds. However, our understanding of the underlying genetics influencing amino acid content and composition remains limited, with only a few candidate genes and quantitative trait loci identified to date. Improved knowledge of the genetics and biological processes that determine amino acid levels will enable researchers to use this information for plant breeding and biological discovery. Towards this goal, we used genomic prediction to identify biological processes that are associated with, and therefore potentially influence, free amino acid (FAA) composition in seeds of the model plant *Arabidopsis thaliana*. Markers were split into categories based on metabolic pathway annotations and fit using a genomic partitioning model to evaluate the influence of each pathway on heritability explained, model fit, and predictive ability. Selected pathways included processes known to influence FAA composition, albeit to an unknown degree, and spanned four categories: amino acid, core, specialized, and protein metabolism. Using this approach, we identified associations for pathways containing known variants for FAA traits, in addition to finding new trait-pathway associations. Markers related to amino acid metabolism, which are directly involved in the FAA regulation, improved predictive ability for branched chain amino acids and histidine. The use of genomic partitioning also revealed patterns across biochemical families, in which serine-derived FAAs were associated with protein related annotations and aromatic FAAs were associated with specialized metabolic pathways. Taken together, these findings provide evidence that genomic partitioning is a viable strategy to uncover the relative contributions of biological processes to FAA traits in seeds, offering a promising framework to guide hypothesis testing and narrow the search space for candidate genes.

## INTRODUCTION

Amino acids play a central role in plant growth and development, as well as human and animal nutrition. In addition to serving as the building blocks for proteins, amino acids are involved in essential biological processes that include nitrogen assimilation, specialized metabolism, osmotic adjustment, alternative energy, and signaling (Rai 2002; Araújo *et al.* 2010; Angelovici *et al.* 2010, 2011; Wu and Messing 2014). Therefore it is no surprise that the homeostasis for absolute levels and relative composition of the free amino acid (FAA) pool is complex and depends, at least in part, on various factors such as allosteric regulation, feedback loops of key metabolic enzymes in amino acid synthesis pathways, and the rate of amino acid degradation (Less and Galili 2008; Jander and Joshi 2010; Hildebrandt *et al.* 2015; Huang and Jander 2017; Amir *et al.* 2018). Studies have also demonstrated that core metabolism has a significant impact on FAA homeostasis. For example, alteration of the interconversion of pyruvate and malate in tomato fruits caused a reduction in FAAs related to aspartate (Osorio *et al.* 2013). In addition, processes related to protein and specialized metabolism also influence FAA homeostasis, especially in vegetative tissues (Hildebrandt *et al.* 2015; Barros *et al.* 2017; Huang and Jander 2017; Hirota *et al.* 2018; Hildebrandt 2018). Together, these lines of evidence highlight that FAA homeostasis is likely determined by orchestration of multiple processes. However, the relative contribution of each process to FAA composition remains unclear.

The composition of the FAA pool is especially critical in dry seeds, since it ensures proper desiccation, longevity, germination, and seed vigor (Angelovici *et al.* 2011; Galili *et al.* 2014). Despite this, very little is known about the function, genetic architecture and regulation of FAAs. The FAA pool comprises 1-10% of total seed amino acid content in maize (Muehlbauer *et al.* 1994; Amir *et al.* 2018) and ∼7% in *Arabidopsis thaliana* (Cohen *et al.* 2014; Amir *et al.* 2018). Although the relative size of the FAA pool is small, manipulation of FAAs in seeds can have a substantial contribution to crop seed nutritional biofortification (Galili and Amir 2013). Nevertheless, several studies have also shown that manipulation of specific FAAs can have pleiotropic effects on growth and germination, indicating their metabolism is intertwined with other key metabolic processes in the seeds (Galili and Amir 2013; Amir *et al.* 2018). Thus, uncovering more about the relative influence of these metabolic processes can help tailor a more effective approach to FAA manipulation and biofortification.

Like many other primary metabolites in dry seeds, FAAs are complex traits with extensive variability and high heritability across natural populations. Multiple genome-wide association (GWA) studies have identified several candidate loci for amino acid traits, both independently (Riedelsheimer *et al.* 2012) and in conjunction with QTL studies (Angelovici *et al.* 2013, 2016). However, the number and effect size of loci detected so far explained only a fraction of the observed phenotypic variation for amino acid traits, with some traits proving harder to dissect than others. For example, (Angelovici *et al.* 2013, 2016) found the strongest associations for traits related to histidine and branched-chain amino acids (BCAAs), but weak signals for most other FAA traits. The findings that amino acid traits frequently have several associated loci and that these loci explain a small proportion of the genetic variation suggest a highly polygenic architecture with many loci of small effect (Korte and Farlow 2013). Additional evidence for metabolic traits indicates that, although many genetic markers may contribute to overall genetic variation, many of these markers are preferentially located in genes that are connected to a biological pathway(s) (Lango Allen *et al.* 2010).

While linkage mapping and GWA studies are typically underpowered to identify variants that are rare and/or of small effect, genomic prediction methods perform well when traits are highly complex (Meuwissen *et al.* 2001; Goddard *et al.* 2009; de Los Campos *et al.* 2013). Genomic prediction models are trained on a subset of individuals with genotypic and phenotypic data, enabling researchers to predict breeding values for genotyped individuals that have an unknown phenotype (Meuwissen *et al.* 2001; Heffner *et al.* 2009). Since its development nearly two decades ago (Meuwissen *et al.* 2001), genomic prediction has dramatically altered the speed and scale of applied genetic and breeding research (Daetwyler *et al.* 2013). The efficacy of genomic prediction results from its simultaneous use of all genotyped markers and indifference to the statistical significance of individual markers, in contrast to analyzing markers one-at-a-time for significance as is done for linkage mapping and GWA studies (Heffner *et al.* 2009). This allows the inclusion of information from all loci to make predictions, instead of basing conclusions only on loci that achieve genome-wide significance, and therefore captures more of the additive genetic variance.

One of the most widely used methods for prediction of complex traits is genomic best linear unbiased prediction (GBLUP) (Meuwissen *et al.* 2001), which assumes that all variants share a common effect size distribution. Recent extensions of the GBLUP model, such as MultiBLUP (Speed and Balding 2014), genomic feature BLUP (Edwards *et al.* 2015, 2016; Sarup *et al.* 2016; Fang *et al.* 2017), and BayesRC (MacLeod *et al.* 2016), incorporate genomic partitions as multiple random effects, allowing effect size weightings to vary across different categories of variants. These partitions can be derived from prior biological information, such as physical position, genic/nongenic regions, pathway annotations, and gene ontologies. Further, genomic partitioning is most successful when a given partition is enriched for causal variant(s) (Sarup *et al.* 2016), providing a framework for guided hypothesis testing. To this end, models that incorporate genomic partitioning have enabled researchers to determine the relative influence of genomic features (e.g. chromosome segments, exons) and/or biological pathways on the variance explained for complex traits. For example, annotations for several biological pathways were used to determine which pathways were associated with udder health and milk production in dairy cattle (Edwards *et al.* 2015). Similarly, gene ontology categories were leveraged to explore the genetic basis of different phenotypes in *Drosophila melanogaster* (Edwards *et al.* 2016). In maize, applications of genomic partitioning models have revealed that SNPs located in exons explain a larger proportion of phenotypic variance compared to other annotation categories (Li *et al.* 2012) and that genomic prediction is improved for multiple traits by incorporating information from gene annotations, chromatin openness, recombination rate, and evolutionary features (Ramstein *et al.* 2020). The inclusion of prior biological information from transcriptomics, GWA studies, and genes identified *in silico* also improved predictions of root phenotypes in cassava (Lozano *et al.* 2017).

In this study, we evaluated genomic partitioning as a method to estimate the relative contribution of metabolic pathway annotations to variation for FAA traits in dry seeds of *Arabidopsis thaliana.* This approach enabled us to incorporate prior knowledge of FAA biochemistry based on metabolic pathways and to identify annotation categories with a disproportionate contribution to the genomic heritability of FAA content and composition. The ultimate objective of this work was to discover metabolic pathway annotations that explained significant variation and improved predictive ability, with the underlying assumption that the corresponding genomic regions are important for determining seed metabolic associations and constraints. These findings can then be considered in future designs to support seed amino acid biofortification.

## MATERIALS AND METHODS

### Plant materials and trait data

For this study, we reanalyzed data of the absolute levels (nmol/mg seed), relative compositions, and biochemical ratios for FAAs in dry *Arabidopsis thaliana* seeds (see **Table S1** for a list of traits). These traits were previously measured by (Angelovici *et al.* 2013, 2016) for 313 accessions of the Regional Association Mapping panel (Nordborg *et al.* 2005; Platt *et al.* 2010). Seeds were obtained from the *Arabidopsis* Biological Resource Center (ABRC, https://abrc.osu.edu/, see **Table S2** for stock numbers). The panel was grown in three independent replicates, each at 18°C to 21°C (night/day) under long day conditions (16 h of light/8 h of dark). Following the desiccation period, dry seeds were harvested and stored in a desiccator at room temperature for at least six weeks prior to analysis to ensure full desiccation (Angelovici *et al.* 2013).

Absolute levels of FAAs (nmol/mg seed) were quantified using liquid chromatography– tandem mass spectrometry multiple reaction monitoring (LC-MS/MS MRM; see Angelovici *et al.* 2013, 2016 for further details). Eighteen of the 20 proteinogenic amino acids were measured, including composite phenotypes for the sum of all FAAs measured (total FAAs) and for each of five biochemical families as determined by metabolic precursor (**Figure S1, Table S1**). This prior knowledge of biochemical relationships among FAAs was also used to determine metabolic ratios, which can represent, for example, the proportion of a metabolite to a related biochemical family or the ratio between two metabolites that share a metabolic precursor (Sauer *et al.* 1999; Weckwerth *et al.* 2004; Wentzell *et al.* 2007). The inclusion of metabolic ratios was based on evidence from multiple studies, which reported novel or more significant associations when using metabolic ratios as compared to absolute levels of metabolites (Wentzell *et al.* 2007; Harjes *et al.* 2008; Vallabhaneni and Wurtzel 2009; Wurtzel *et al.* 2012; Lipka *et al.* 2013; Gonzalez-Jorge *et al.* 2013; Angelovici *et al.* 2013, 2016; Owens *et al.* 2014).

For each amino acid, relative composition was calculated as the absolute level over the total. Additional ratio traits were determined based on biochemical family affiliation (Angelovici *et al.* 2016). Traits and their respective abbreviations are described in **Table S1**. Overall, the 65 traits included 25 absolute FAA levels (individual amino acids and composite traits), 17 relative levels (ratio of the absolute level for an amino acid compared to total FAA content), and 23 family-derived traits (ratio of the absolute level for an amino acid to the total FAA content within a given family).

Following the guidelines for multi-stage genomic prediction (Piepho *et al.* 2012), the best linear unbiased estimates (BLUEs) for each accession were used as the phenotypic data in this study and were calculated using the HAPPI-GWAS pipeline (Slaten *et al.* 2020a) in R v3.6.0 (R Core Team 2016). First, outlier removal was performed by fitting a mixed effects model using the ‘lmer’ function in the ‘lme4’ package (v1.1-21, (Bates *et al.* 2015), with the raw trait values as the response variable, replicate included as a random effect, and accession included as a fixed effect. Studentized deleted residuals were then used to identify outliers (Kutner *et al.* 2004). Following outlier removal, the Box-Cox transformation (Box and Cox 1964) was applied for each trait to avoid violating model assumptions of normally distributed error terms and constant variance. Finally, to remove phenotypic variability arising from environmental conditions, the BLUE for each accession was obtained from the fitted mixed model described above, which was applied across all three replicates. The BLUEs for each trait were used as the response variables in all subsequent prediction models.

### Genetic data

The accessions used in this study were previously genotyped using a 250k SNP panel (v3.06, Atwell *et al.* 2010). The software PLINK (v1.9, Purcell *et al.* 2007) was used to filter for minor allele frequency (MAF) > 0.05, reducing the number of SNPs from 214,051 to 199,452. Principal component analysis was performed on this filtered SNP set using the ‘prcomp’ function in R. The first two principal components explained 5.6% of the variance (**Figure S2**) and were included as fixed covariates in the prediction models.

### Selection of pathway SNPs

To examine specific metabolic pathways, SNPs were selected based on annotation categories from the MapMan annotation software (Thimm *et al.* 2004) for the TAIR10 version of *Arabidopsis* (Berardini *et al.* 2015). We focused broadly on 20 pathways, which spanned four categories: amino acid metabolism (three pathways), core metabolism (three pathways), specialized metabolism (five pathways), and protein metabolism (nine pathways) (**Table 1**), all of which are known to be involved in FAA metabolism to some extent. The SNP positions were first matched to the corresponding Ensembl gene id using the ‘biomaRt’ package (Durinck *et al.* 2005, 2009) in R. We then selected all SNPs within a 2.5 kb range of the start and stop position for each gene, which is within the range of the estimated average intergenic distance in *Arabidopsis* (Zhan *et al.* 2006) and includes upstream promoter regions. Relative SNP positions for each pathway are provided in **Figure S3**. Pathways and MapMan annotation categories, including the number of genes and SNPs represented, are described in **Table 1**. We used MapMan annotations for all genes except BCAT2 (At1g10070), which was moved from the amino acid synthesis pathway to the amino acid degradation pathway along with other SNPs in the same haploblock (chromosome 1, 3274080 to 397645 bp). This decision was based on previous work, which showed that *bcat2* mutants accumulate higher levels of branched-chain amino acids in seeds, thereby demonstrating that BCAT2 has catabolic activity (Angelovici *et al.* 2013).

**Table 1.** Summary of selected biological pathways.

| Pathway | Number of genes<sup>a</sup> | Number of SNPs<sup>a</sup> | MapMan BINCODE |
| --- | --- | --- | --- |
| ***Amino Acid Metabolism*** |  |  |  |
| amino acid synthesis | 376 | 2084 | 13.1 |
| amino acid degradation | 160 | 1094 | 13.2 |
| amino acid transport | 144 | 939 | 34.3 |
| ***Primary Metabolism*** |  |  |  |
| glycolysis | 148 | 858 | 4 |
| TCA cycle | 167 | 926 | 8 |
| ATP synthesis (alternative oxidase) | 10 | 66 | 9.4 |
| ***Specialized Metabolism*** |  |  |  |
| isoprenoids | 269 | 1788 | 16.1 |
| phenylpropanoids | 161 | 845 | 16.2 |
| nitrogen containing | 39 | 229 | 16.4 |
| sulfur containing | 113 | 733 | 16.5 |
| flavonoids | 171 | 1062 | 16.8 |
| ***Protein Metabolism*** |  |  |  |
| amino acid activation | 203 | 1231 | 29.1 |
| protein synthesis | 1383 | 7290 | 29.2 |
| protein targeting | 624 | 3689 | 29.3 |
| protein posttranslational modification | 1407 | 8794 | 29.4 |
| protein degradation | 996 | 6405 | 29.5 |
| ubiquitin | 2691 | 16000 | 29.5.11 |
| protein folding | 138 | 814 | 29.6 |
| protein glycolysis | 87 | 459 | 29.7 |
| protein assembly | 44 | 312 | 29.8 |

<sup>a</sup>Includes a 2.5 kb buffer before and after the start/stop position of each gene

### Prediction models

The Linkage Disequilibrium Adjusted Kinship (LDAK) software (v5.0, Speed *et al.* 2012, http://dougspeed.com/ldak/) was used to implement two models for genomic prediction of each trait: GBLUP, which uses a single marker-based additive genetic relatedness matrix, and MultiBLUP, which incorporates multiple marker-based additive genetic relatedness matrices, each calculated with different subsets of genome-wide markers (Speed and Balding 2014).

Genomic prediction was performed for all markers (*p* = 199,452) using a GBLUP model, in which individuals were included as a random effect and the additive genetic relatedness matrix was used as part of the variance-covariance matrix among the individuals (Whittaker *et al.* 2000; Meuwissen *et al.* 2001). First, the pairwise genetic similarity between individuals was estimated using a genomic similarity matrix (GSM), or kinship matrix (VanRaden 2008; Astle and Balding 2009):

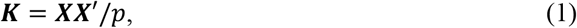

where *X* is an *n x p* design matrix of SNP genotypes, *X*′ is the transpose of *X, n* is the total number of individuals, and *p* is the total number of markers. The GBLUP model was then fit with the random effects model:

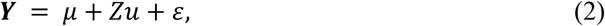

where *Y* is the vector of phenotypic values for *n* individuals, *μ* is the overall mean, *Z* is a design matrix connecting observations to genotypes, *u* is the vector of random genetic effects distributed as 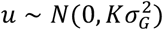, and *ε* is the random error term distributed as 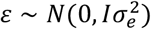, where *K* is the GSM representing the correlation structure of *u, I* is a *n* × *n* identity matrix, and 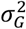 and 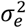 are variances.

The MultiBLUP model (Speed and Balding 2014) was used to incorporate biological pathway information into genomic prediction. As an extension of the GBLUP model, MultiBLUP is a multi-kernel model that subdivides genetic effects into at least two random effects, where different subsets of markers are used to calculate GSMs for each random effect. In this study, the MultiBLUP model included a random genetic effect corresponding to sets of markers within a single biological pathway (*m*) and a second random genetic effect corresponding to the remaining markers not included in the given pathway (∉ *m*). Using notation from equation (2) and Speed and Balding (2014), the MultiBLUP model within the context of this work is:

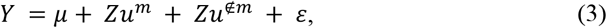

where *u*^*m*^ is the vector of random genetic effects distributed as 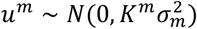, with *K*^*m*^ representing the kinship matrix calculated using markers within a given biological pathway and 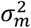 denoting the corresponding variance component; *u*^∉*m*^ is the vector of random genetic effects distributed as 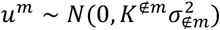, with *K*^∉*m*^representing the kinship matrix calculated using markers outside of the given biological pathway and 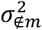 denoting the corresponding variance component; and ***Y***, Z, *μ*, and *ε* are as previously described.

For our purposes, kinship matrices were estimated using the LDAK software for either all SNPs (GBLUP) or each SNP partition (MultiBLUP, i.e. separately for SNPs belonging to a single pathway and all other remaining SNPs). For each pathway, the extent of collinearity due to LD was determined by examining Spearman’s rank correlation between the off-diagonal elements of the kinship matrices for the pathway SNPs and remaining genomic SNPs. For estimates of genomic heritability, values were constrained to be positive and less than one. Additionally, the parameter α, which models the relationship between heritability and MAF, was set to α = 0 under the assumption that SNPs with lower MAF contribute less to heritability than SNPs with higher MAF (Speed *et al.* 2017). In relation to Eq. 2 and 3, the parameter α adjusts the expected contribution of each SNP to heritability, with a value of α = −1 assuming that heritability does not depend on MAF (see Speed et al. 2017 for details).

### Estimation of genomic heritability

For both GBLUP and MultiBLUP, average information restricted maximum likelihood (REML, see Speed and Balding 2014 for details) was used to compute variance component estimates for 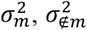, and 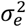. The maximum number of iterations to achieve convergence was set to 500. This process was repeated for each trait and pathway combination. In the case of the GBLUP model, 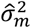 is the estimate of variance for all SNPs. These estimates were used to calculate genomic heritability as the ratio of additive genomic variance explained for a given marker set 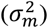 over the total variance explained (the sum of 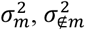, and the residual variance, 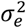):

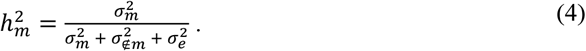

For the MultiBLUP model, the proportion of genomic heritability explained was calculated as:

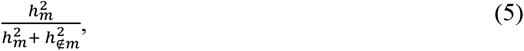

where 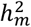 is the genomic heritability explained by SNPs in a given genomic partition and 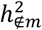 is the genomic heritability explained by all other SNPs not included in the partition.

### Assessing model performance

The performance of the prediction models was determined using ten-fold cross validation with a one-fold holdout, with the same training and testing sets used for both the GBLUP and MultiBLUP models. For each cross validation, the genomic estimated breeding value (GEBV) was derived from marker data for the excluded individuals based on estimates of random genetic effects for the individuals in the training set. This process was repeated five times for a total of 50 cross validations per trait and pathway combination. Predictive ability was then calculated as *r*(*ĝ, g*), where *ĝ* represents the GEBVs and *g* represents the BLUEs for each trait. Reliability, which is the coefficient of determination (*r*^2^) scaled by heritability, was calculated as 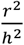 (Rincent *et al.* 2012). Bias was calculated as the simple linear regression slope estimate between the GEBVs and BLUEs for each trait, with a slope estimate of one indicating no bias. Lastly, the overall root mean squared error (RMSE), which measures prediction bias and variability, was calculated as the square root of the mean for the squared difference between the BLUEs and GEBVs across all cross-validations, 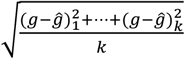, where *k* is the number of cross-validations. Predictive abilities for the MultiBLUP and GBLUP models were compared using a one-sided, paired Welch’s t-test for unequal variances.

### Generation of an empirical null distribution

To test if a metabolic pathway explained more variation than expected by chance, we generated an empirical null distribution for each trait and pathway combination. The null hypothesis was that a given biological pathway will explain a similar amount of trait variance as the same number of SNPs in randomly selected gene groups (Edwards *et al.* 2015). To establish a null distribution, we first defined 1000 random gene groups for each pathway, where the target number of SNPs in each random gene group was comparable to the size of the pathway. Ranges for the number of genes and SNPs sampled for each pathway are provided in **Table S3**. For each random subset, all SNPs within 2.5 kb of the start and stop positions were sampled for a randomly selected gene. This process was repeated by randomly sampling genes one at a time until the target number of SNPs for each subset was achieved. Genes within a given pathway were excluded from the random sampling procedure for that pathway. As discussed in Edwards *et al.* (2015), this approach does not explicitly model variation in other parameters (e.g., allele frequencies, LD), but it is expected that these differences are captured to some extent by the sampling process.

Next, we used two metrics to test if SNPs in a given pathway explained more genomic variance than expected by chance and increased model fit for each trait: (1) the proportion of genomic heritability explained by a pathway compared to the random gene groups described above, and (2) the likelihood ratio (LR) test statistic as a measure of pathway model fit compared to the model fit of random gene groups. The proportion of heritability explained was calculated as described previously in equation (5) and the LR test statistic was calculated as twice the difference between the log likelihood of the MultiBLUP model and the log likelihood of the GBLUP model. For each pathway and trait combination, the values for proportion of heritability explained and the LR test statistic were compared to the empirical cumulative distribution function for the corresponding 1000 random gene groups using the ‘ecdf’ function in R. To determine if the observed value was greater than the random values for each metric, *P*-values were computed with a one-sided test using the ‘t_test’ function in the R package ‘rstatix’ (Kassambara 2020).

### Correction for multiple testing

For each trait, the Benjamini-Hochberg procedure (Benjamini and Hochberg 1995) was used to adjust for multiple testing across pathways (n = 20) at a 10% false discovery rate (FDR). Multiple testing correction was performed with the ‘p.adjust’ function in R for the proportion of heritability explained, the LR test statistic, and predictive ability.

### Identifying biological pathways of interest

In summary, a pathway was considered of interest for a trait if the MultiBLUP model passed all three of the following criteria:

1. The proportion of heritability explained was significantly greater than empirical values for random gene groups of the same size (FDR-adjusted *P*-value < .10),
2. The LR test statistic was significantly greater than empirical values for random gene groups of the same size (FDR-adjusted *P*-value < .10),
3. The MultiBLUP model significantly improved predictive ability compared to the GBLUP model (FDR-adjusted *P*-value < .10).

Together, criteria (1) and (2) established that a given pathway improved model fit better than a random set of SNPs. Criteria (3) was imposed to ensure that there was a meaningful difference in predictive ability when pathway information was incorporated via MultiBLUP compared to the naive GBLUP model that incorporated no pathway information.

### Data availability statement

Genotype data were previously published (Atwell *et al.* 2010) and were accessed from github.com/Gregor-Mendel-Institute/atpolydb/wiki. The scripts and phenotypic data supporting the conclusions of this article are publicly available as a Snakemake workflow (v5.4.2, Köster and Rahmann 2012) on GitHub at github.com/mishaploid/aa-genomicprediction. **Table S2** details the ABRC stock names and accession numbers for each individual.

## RESULTS AND DISCUSSION

In this study, we applied a genomic partitioning model to evaluate the contribution of metabolic pathways to FAA traits in seeds. The combination of a genomic partitioning framework and the model system *Arabidopsis* allowed us both to test the feasibility of this approach and to further examine the relative contribution of each pathway to the genetic basis of FAA traits in seeds. Additionally, because FAA traits are part of core metabolism that is highly conserved, we hypothesize that our findings can be used to develop hypotheses in crop systems, where there is potential to contribute to the biofortification of essential amino acids.

### Genomic prediction was most effective for absolute levels of free amino acids

We first established the efficacy of standard GBLUP in a diversity panel of 313 *Arabidopsis* individuals, which represents a substantial proportion of the known genetic variability present in *Arabidopsis* (Nordborg *et al.* 2005). Because this setting is distinct from the closed breeding populations of dairy cattle, maize, and other agricultural species where genomic prediction is often applied (e.g. Heffner *et al.* 2009; Wolc *et al.* 2016; Weller *et al.* 2017), we were interested in testing how well genomic prediction would work in this panel. We were also interested in testing the utility of genomic prediction for FAA traits, which are highly conserved.

Using the GBLUP model, we observed low to moderate predictive ability for the amino acid traits measured (**Table 2**). Of these 65 FAA traits, 30 had a predictive ability greater than 0.3 (**Figure 1, Table 2**). In general, prediction was effective for a greater number of absolute level FAA traits, with 21 out of 25 absolute traits having a predictive ability > 0.3 (84%), compared to relative levels (4 out of 17, 24%) and family-derived ratios (5 out of 23, 22%). The family ratio of methionine (met_AspFam) had the highest predictive ability (r = 0.47), while the relative level of serine (ser_t) had the lowest predictive ability (r = 0.08) (**Table 2**). The observation of moderate prediction accuracies for many of these traits (**Figure 1, Table 2**) suggests that there is linkage disequilibrium (LD) between markers and causal loci, providing evidence that genomic prediction can be successfully applied in this system.

**Table 2.**
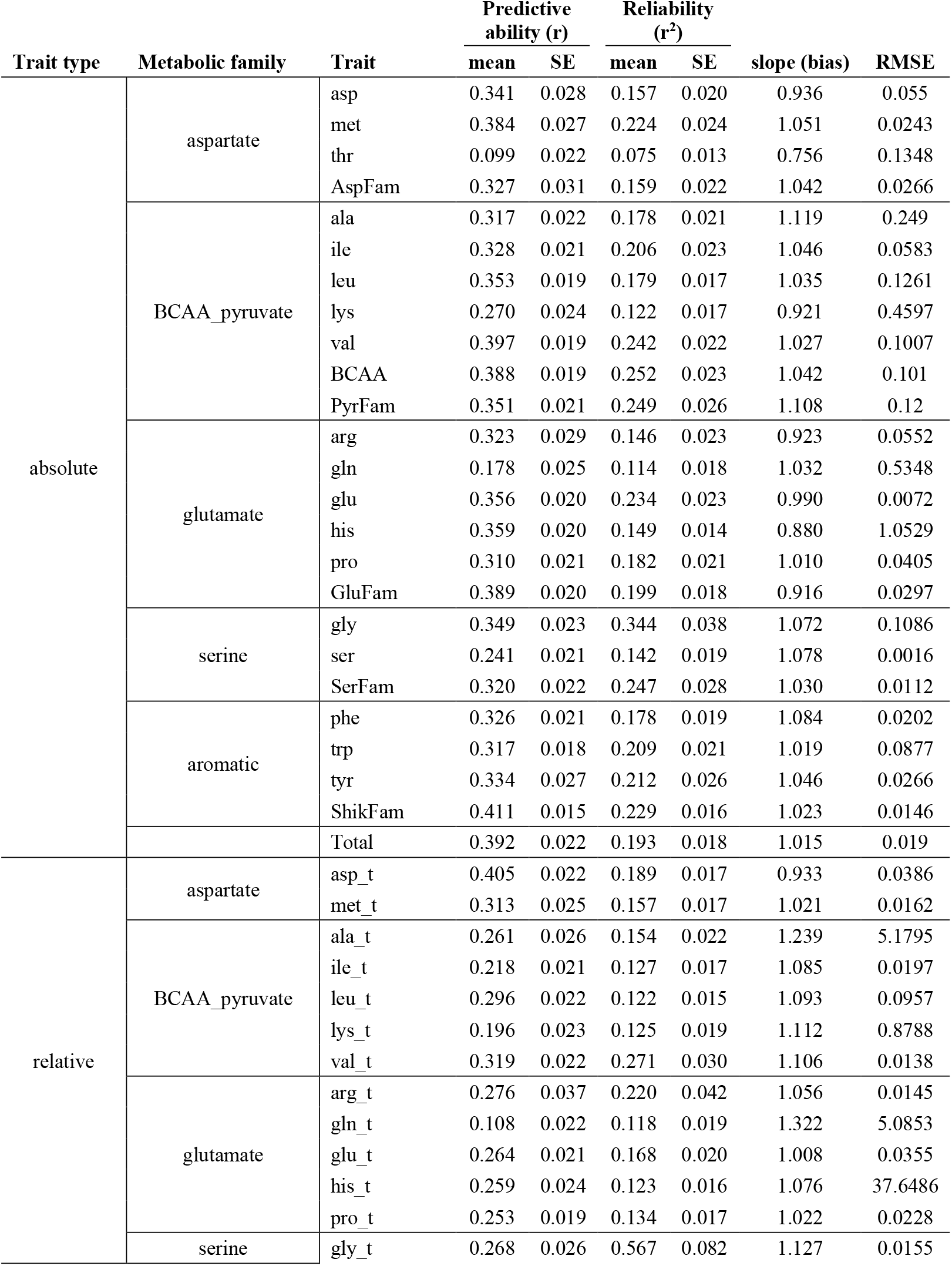
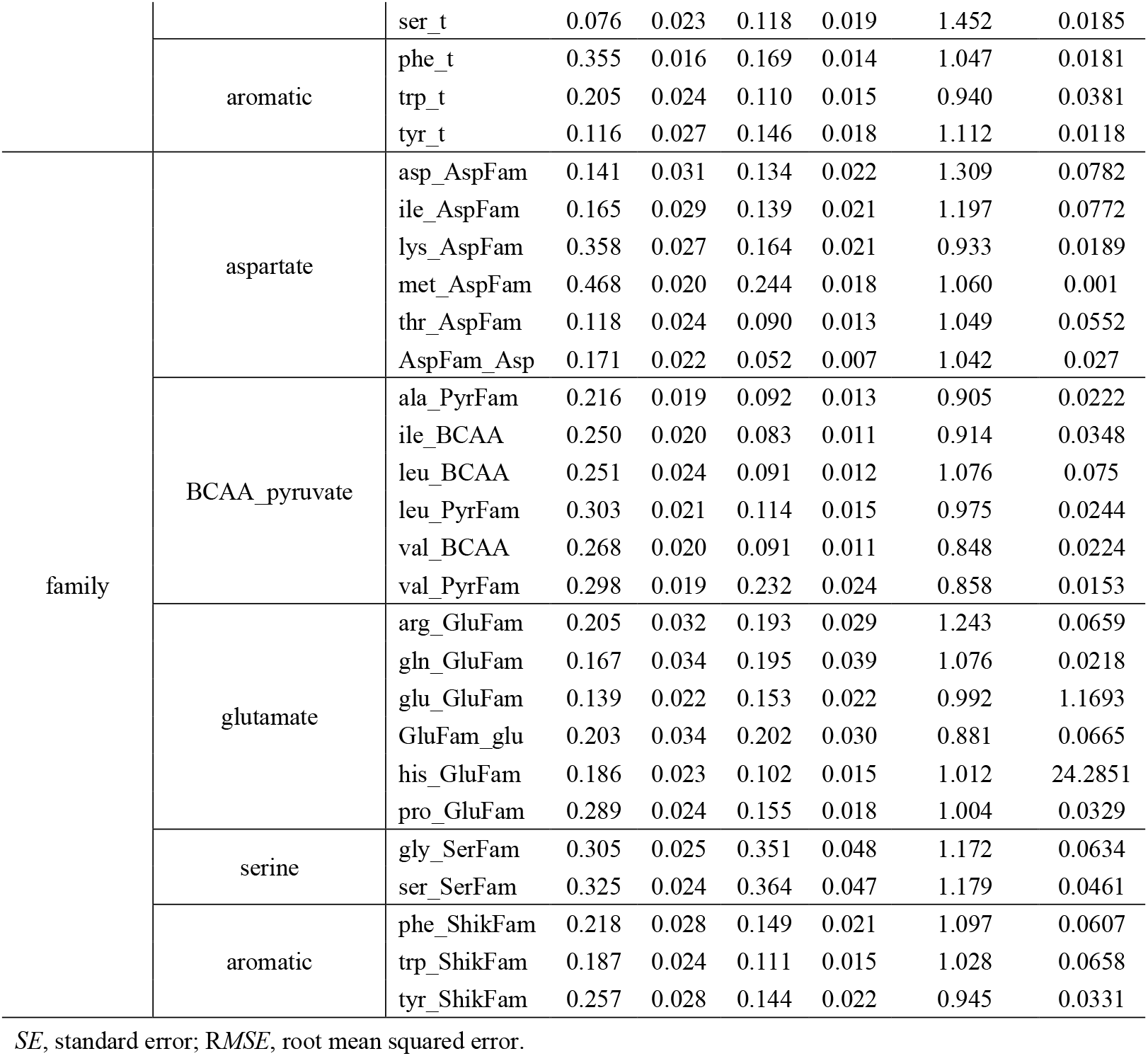
Genomic prediction results for 65 free amino acid traits using a GBLUP model.

**Figure 1.**
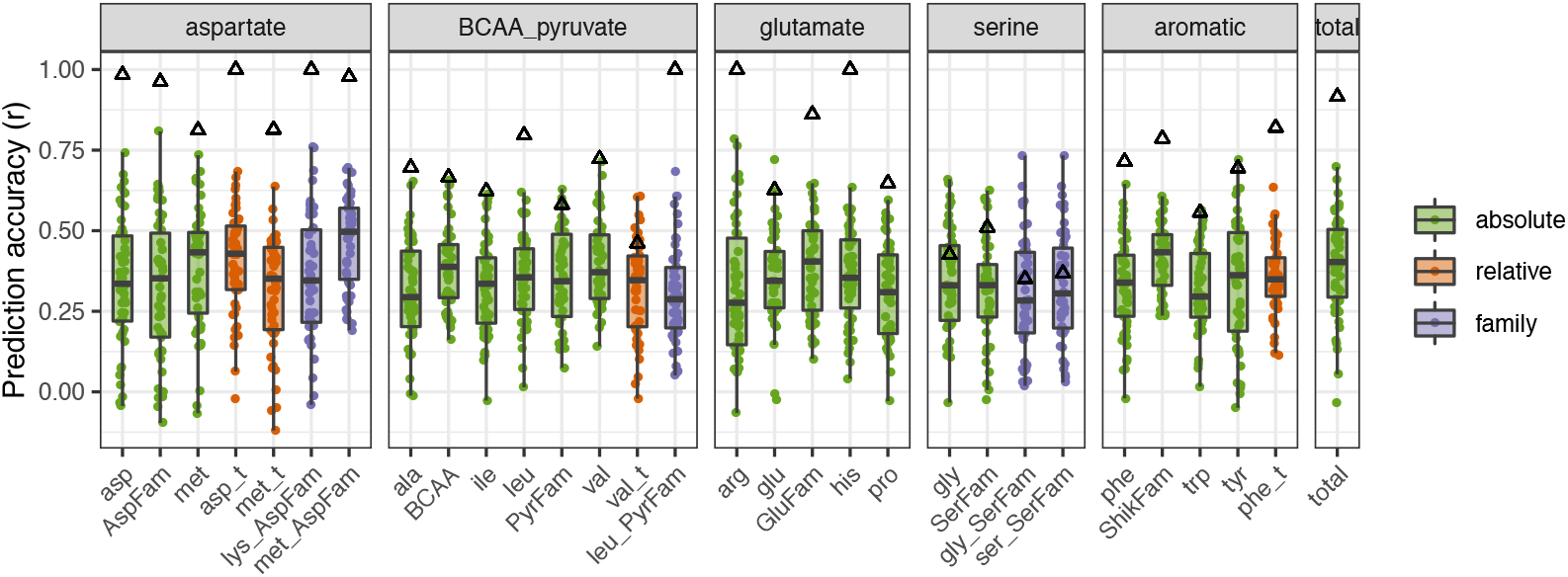
Genomic prediction performed well for a higher proportion of absolute traits compared to relative and family-based ratio traits. Boxplots show free amino acid traits with predictive ability (r) > 0.3 based on genomic best linear unbiased prediction (GBLUP). Black triangles indicate the genomic heritability for each trait. Colors indicate whether the trait is an absolute level, relative level, or family-based ratio. Each point represents an individual cross-validation.

### Annotations for biological pathways explained significant variation and improved predictive ability of free amino acid traits in seeds

We next applied a genomic partitioning approach, MultiBLUP, to investigate the association of different metabolic annotation categories with FAA traits in dry *Arabidopsis* seeds. The focus was specifically on categories which are thought to influence FAA homeostasis, but where the degree of this influence is unclear, especially in dry seeds (Skirycz et al. 2010, 2011; Hildebrandt et al. 2015; Hildebrandt 2018).

The pathway annotations listed in **Table 1** were used to subset SNPs and spanned the broad categories of amino acid, core, specialized, and protein metabolism. When partitioning these pathways in the MultiBLUP model, 18 trait-pathway combinations were flagged as potentially related based on comparison to a null distribution (**Figure 2A, Table 3**). The observation that specific pathways improved model fit based on the LR test statistic, explained a significant proportion of genomic heritability, and improved predictive ability suggests that these pathway annotations may have biological relevance for FAA traits.

**Table 3.**
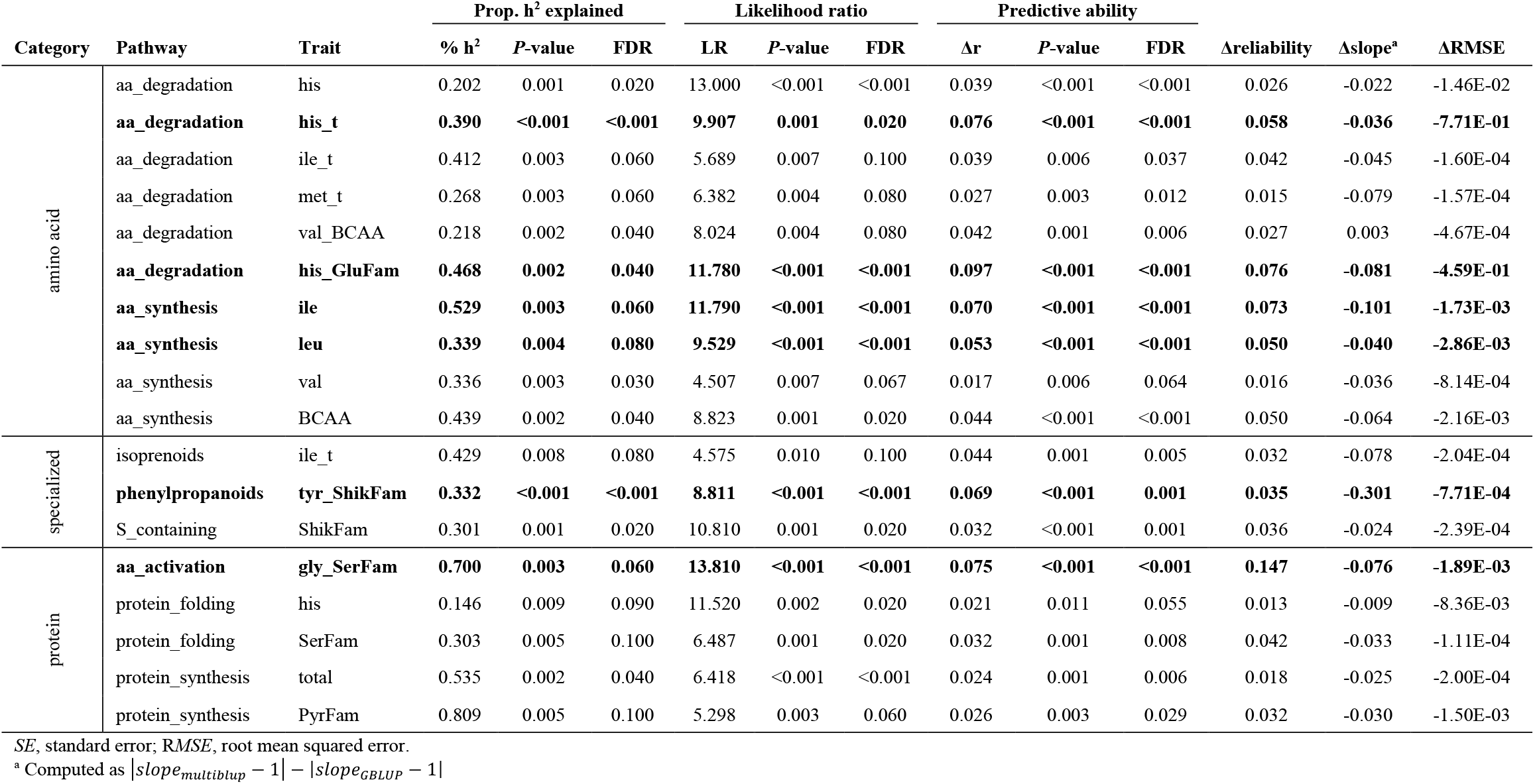
Free amino acid traits and pathway combinations for which the MultiBLUP model explains a significant proportion of heritability, improves model fit relative to random gene groups of approximately the same size, and increases accuracy compared to GBLUP.

**Figure 2.**
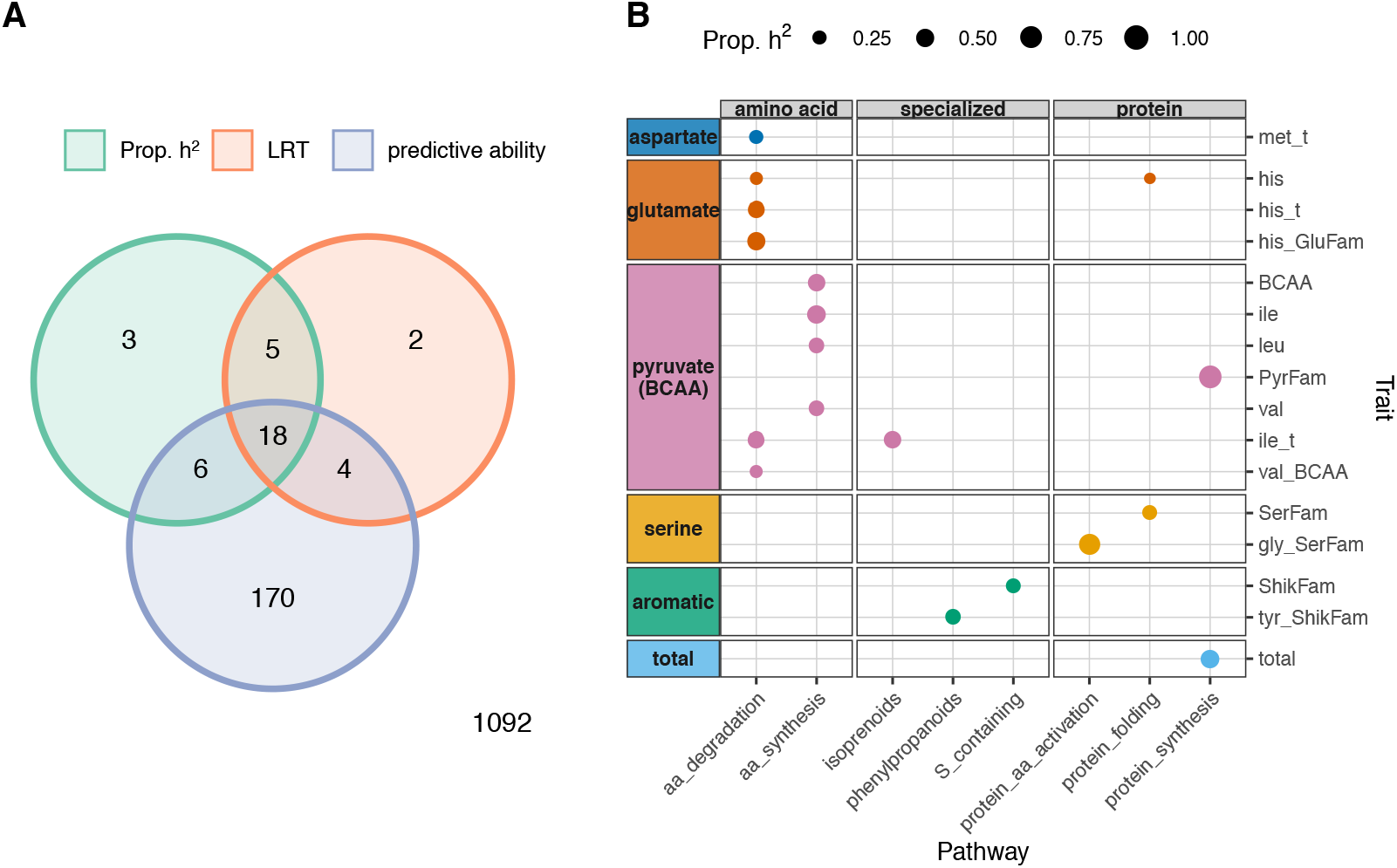
Biological pathways explain significant variation and improve predictive ability for free amino acid traits when incorporated into a MultiBLUP model. **(A)** Venn diagram showing which trait-pathway combinations passed significance criteria (FDR adjusted *P*-value < .10) for proportion of heritability explained (Prop. h^2^), likelihood ratio test statistic (LRT), and improved predictive ability for MultiBLUP compared to GBLUP. The bottom right corner indicates the number of combinations that did not pass any significance criteria. The Venn diagram was constructed using the ‘limma’ package in R (Smyth *et al.* 2005). **(B)** Dots indicate trait-pathway combinations that passed all three significance criteria. The diameter of each dot is proportional to the amount of genomic variance explained by pathway SNPs in the MultiBLUP model. Traits are included on the y-axis and are grouped by metabolic family (aspartate, glutamate, pyruvate/BCAA, serine, aromatic). Pathways are included on the x-axis and separated into amino acid, core, specialized, and protein metabolism categories.

For the trait-pathway combinations that passed the significance criteria, the MultiBLUP model generally reduced bias and RMSE compared to the GBLUP model (**Table 3**). For six of these trait-pathway combinations, the predictive ability for the MultiBLUP model was also over 5% higher than for the GBLUP model (**Table 3**, bold). This substantial increase in predictive ability was observed in the pyruvate/BCAA family for absolute levels of leucine (leu, 5.3%) and isoleucine (ile, 7%) when the model included SNPs in the amino acid synthesis pathway. The highest increase in predictive ability was observed when incorporating the amino acid degradation pathway for traits in the glutamate family, which included the relative level and family-based ratio for histidine (his_t, 7.6%; his_GluFam, 9.7%). A similar increase in predictive ability was observed when including SNPs related to phenylpropanoids for the family ratio of tyrosine (tyr_ShikFam, 6.9%) and when including SNPs related to protein amino acid activation for the family ratio of glycine (gly_SerFam, 7.5%).

### Amino acid synthesis and degradation pathways were significantly associated with several FAA traits

The homeostasis of FAAs is regulated by multiple allosteric enzymes and feedback loops (Less and Galili 2008; Jander and Joshi 2010; Hildebrandt *et al.* 2015; Huang and Jander 2017; Amir *et al.* 2018). However, the homeostasis of some FAAs, such as proline, can also be determined by environmental conditions. For example, proline may serve as either an osmoprotectant under stress or an energy source during development, and its elevation is mostly from active synthesis (Szabados and Savouré 2010; Hayat *et al.* 2012). In addition, previous work has suggested that an overarching metabolic switch occurs during late maturation to desiccation, when amino acid synthesis is active (Fait *et al.* 2006). Hence, our initial hypothesis was that FAA traits would be strongly associated with pathway annotations within core and amino acid metabolism.

For amino acid metabolism, our initial hypothesis was supported by significant associations between the amino acid degradation pathway and with six traits, which spanned the aspartate, glutamate, and pyruvate/BCAA families (**Figure 2B**). Two BCAA traits, ile_t and val_BCAA, were associated with amino acid degradation, consistent with previous work which identified a large effect QTL that explained 12-19% of the variance for BCAA traits (Angelovici *et al.* 2013). Based on this previous work, the causal gene was identified as the catabolic branched-chain amino acid transferase 2 (BCAT2; At1g10070) (Angelovici *et al.* 2013). Our results recapitulate this finding, showing that the amino acid degradation pathway, which contains the BCAT2 haploblock, explained both a significant proportion of heritability (41%) and improved predictive ability for BCAA traits (e.g. by 3.9% for ile_t) (**Table 3**). In contrast, the only additional associations that were identified were between amino acid synthesis and BCAA traits, despite no prior evidence that QTLs for BCAAs contain genes related to amino acid synthesis (Angelovici *et al.* 2013). Surprisingly, these were also the only associations that were identified for amino acids synthesis, despite evidence that levels of several FAAs and transcription of their biosynthetic genes are elevated toward desiccation (Fait *et al.* 2006). This observation could arise from one of several reasons: 1) the elevation of transcription for amino acid biosynthetic genes does not lead to a corresponding elevation in metabolic pathway products, 2) our sample size and statistical approach was unable to resolve other traits associated with amino acid synthesis, or 3) we are unable to cleanly partition pathway SNPs from background genome wide markers. Nonetheless, our results imply that amino acids synthesis may be more important for BCAAs than for other FAA traits at this stage of development.

We also observed that annotations for amino acid degradation were associated with histidine and methionine FAA traits (**Figure 2B, Table 3**), which, to our knowledge, has not been reported in previous QTL studies for seed FAAs. Both histidine and methionine are essential amino acids, which are deficient in most crop seeds, and therefore of special interest for biofortification and crop improvement (Galili and Amir 2013). Notably, very little is currently known about the pathway for histidine degradation in plants. Taken together, these findings suggest that the MultiBLUP approach can not only recapture previous observations for FAA traits, but can also generate new insights into their genetic regulation.

Both amino acid and core (or primary) metabolism are tightly interconnected. For example, amino acids in the glutamate family are known to play a central role in core metabolism, mainly by functioning as precursors for energy generation via glycolysis, amino acid metabolism, and the TCA cycle. However, we found no associations for any FAA traits with the core/primary metabolic pathways tested in this study, which included glycolysis, the TCA cycle, and ATP synthesis via alternative oxidase (**Figure 2B, Table 3**).

### Gene annotations for specialized metabolism are associated with FAA precursors

The synthesis of specialized metabolites involves many FAAs. For example, methionine and aromatic amino acids (i.e. phenylalanine, tryptophan, and tyrosine) are precursors for alkaloids, phenylpropanoids, and glucosinolates. Levels of these specialized metabolites are often dependent on the availability of their FAA precursors (Tzin and Galili 2010; Maeda and Dudareva 2012). However, less is known regarding whether the extensive natural variation of these specialized metabolites produces a feedback effect on FAA precursors, especially in seeds. Previous work in vegetative tissues has found that perturbation of the synthesis for secondary metabolites produces a pleiotropic effect on other types of metabolism, including FAAs (Chen *et al.* 2012; Slaten *et al.* 2020b), but the nature of such interactions is not well understood.

Consistent with knowledge of precursors for specialized metabolites, we observed that aromatic FAAs were associated with categories belonging to specialized metabolism (**Figure 2**). This included associations for the combined absolute levels of FAAs in the shikimate family (ShikFam) with the pathway for sulfur-containing compounds and between the family ratio of tyrosine (tyr_ShikFam) with the phenylpropanoid pathway. When partitioning SNPs from the phenylpropanoid pathway in the MultiBLUP model, we observed a 6.9% increase in predictive ability for tyr_ShikFam (**Table 3**), suggesting SNPs in this pathway have a substantial contribution to the variation for Tyr_ShikFam or are in strong LD with one or more causal variants. We also found an unexpected association between isoprenoid metabolism and the relative ratio of isoleucine (ile_t), which is part of the BCAA family (**Figure 2B**). The metabolic relationship is less clear in this case, as isoleucine is not directly involved in phenylpropanoid metabolism, and provides an avenue for further investigation.

A recent metabolic GWA study identified an unanticipated association between glucosinolate biosynthesis and levels of free glutamine in seeds of *Arabidopsis* (Slaten *et al.* 2020b). This finding was further validated by evidence that elimination of seed glucosinolates significantly impacted levels of glutamine during early seed development (Slaten *et al.* 2020b). Notably, when partitioning SNPs for sulfur-related metabolism, the family-based ratio for glutamine (GluFam_glu) passed significance criteria for proportion of heritability explained (40.5%, FDR corrected *P*-value = .10) and predictive ability (3.7% increase compared to GBLUP, FDR corrected *P*-value = .006), but not for the LR test statistic (5.48, FDR corrected *P*-value = .12). This observation reinforces that additional studies, especially with greater statistical power, may identify more connections with biological relevance.

### Annotations for protein metabolism are associated with serine family FAAs

It stands to reason that FAA homeostasis will be influenced by protein metabolism since FAAs serve as the building blocks for proteins. Consistent with this expectation, significant increases in FAAs are observed under many abiotic stresses and suggested to result from protein autophagy and turnover (Hildebrandt *et al.* 2015; Barros *et al.* 2017; Huang and Jander 2017; Hirota *et al.* 2018; Hildebrandt 2018). In contrast, the *opaque2* null mutant in maize exhibits a reduction in the most abundant seed storage proteins and a significant elevation of many FAAs, despite an unchanged composition of protein-bound amino acids (Wang and Larkins 2001; Schmidt *et al.* 2011), indicating a complex relationship between the free and bound amino acid pools for protein metabolism. Hence, it is unclear to what extent protein metabolism affects FAAs, particularly in seeds where protein composition is critical for nutritional quality.

Interestingly, we find that protein metabolism annotations are associated with five FAA traits, which spanned the glutamate, pyruvate/BCAA, and serine families, and included the composite trait for total FAA content (**Figure 2, Table 3**). Notably, no aromatic FAA traits were associated with protein metabolic annotation categories, while the serine family FAA traits were exclusively associated with this group of pathways. Further, the family-based ratio for glycine (Gly_SerFam) showed an increase in predictive ability of 7.5% when partitioning SNPs related to amino acid activation in the MultiBLUP model (**Table 3**). This suggests that genes related to amino acid activation, such as tRNA synthetases, may contribute to the homeostasis of glycine and serine. Overall, even though most protein metabolism occurs at seed maturation, we found evidence that annotations for protein metabolism influence FAAs in dry seeds, suggesting that FAA levels at this stage may reflect prior events occurring earlier in seed development.

### Pathway size influences proportion of heritability explained, model fit, and predictive ability

To examine the relationship between pathway size, LD, and variance partitioning, we compared off-diagonal elements of the kinship matrices for pathway SNPs and remaining genomic SNPs (**Figure 3A**). Spearman’s correlations ranged from 0.17 for the ATP synthesis via alternative oxidase category (e_alt_oxidases, 66 SNPs, 0.03% of total SNPs) to 0.85 for the protein degradation by ubiquitin category (degradation_ubiquitin, 16000 SNPs, 8.02% of total SNPs) (**Figure 3A**). In general, pathways containing a greater number of SNPs displayed more collinearity with SNPs not contained in the pathway. Similar to observations for genomic partitioning based on gene ontology terms for locomotor activity in *Drosophila* (Rohde *et al.* 2018), we observe that pathways which increased predictive ability also explained a large proportion of genomic heritability, whereas pathways with a greater number of SNPs explained less genomic heritability and did not improve predictive ability (**Figure 3B**). Further, as suggested by Rohde *et al.* (2018), pathways which explained all of the genomic heritability likely represent an overestimation caused by high similarity between the relationship matrices for the pathway and background genomic SNPs.

**Figure 3.**
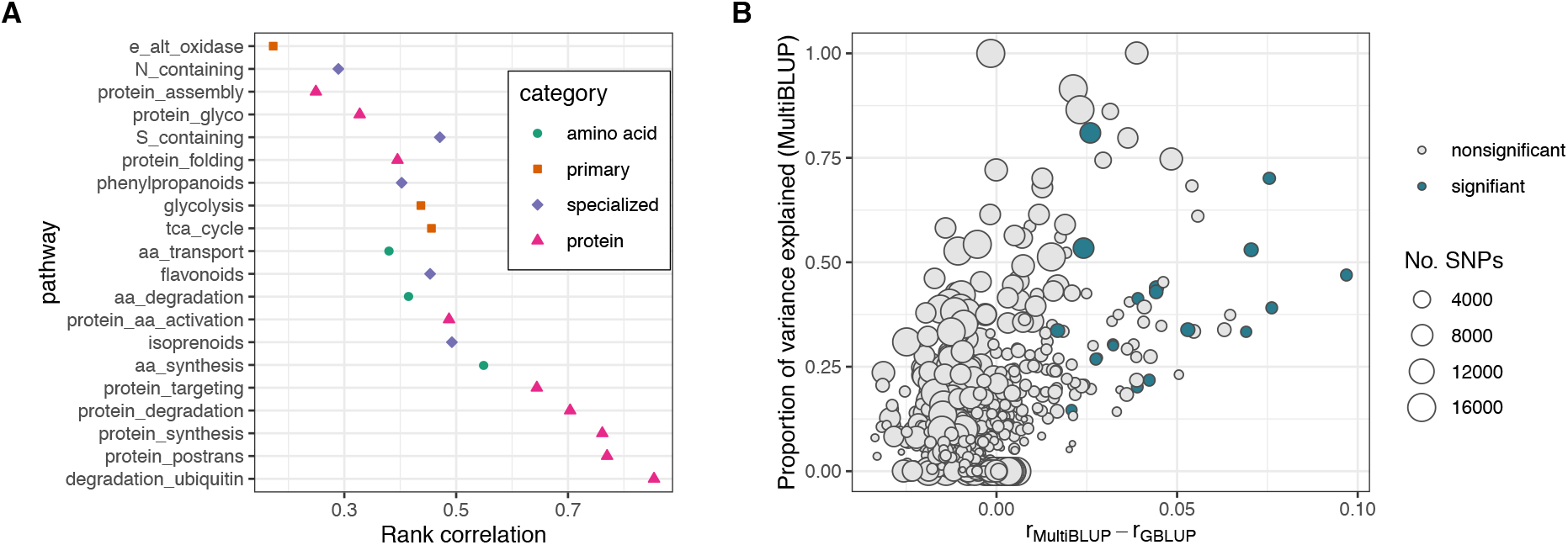
Pathway size influences the proportion of heritability explained and predictive ability when using a MultiBLUP model. **(A)** Spearman’s rank correlations between off-diagonal elements of the kinship matrices for each pathway and the remaining genomic SNPs. Pathways are sorted from top to bottom by increasing size (number of SNPs). **(B)** Difference in predictive ability between the MultiBLUP and GBLUP models compared to the proportion of heritability explained by each pathway for all 1300 trait-pathway combinations (65 traits, 20 pathways). The diameter of the points is proportional to the number of SNPs in the pathway and color indicates whether or not a trait-pathway combination passed significance for proportion of heritability explained, likelihood ratio test statistic, and predictive ability.

## Conclusion

Overall, we find that predictive ability for FAA traits was improved by incorporating prior knowledge from metabolic pathway annotations for several FAA traits, adding to a growing body of literature that demonstrates the utility of genomic partitioning in the study and prediction of complex traits. This study further highlights that specific metabolic pathways are associated with natural variation of FAA traits across amino acid families. The amino acid degradation pathway was significantly associated with traits in the BCAA/pyruvate, glutamate, and aspartate families, while specialized metabolism was associated with traits in the aromatic family and protein metabolism was associated with traits in the serine, pyruvate/BCAA, and glutamate families. Thus, although the FAA metabolic network is tightly connected, the predominant genetic architecture underlying variation for specific FAA traits varies, at least for this stage of seed development. Overall, this study furthers our understanding of the contribution from specific metabolic pathway genes to amino acid trait variation and offers an additional strategy to investigate other complex metabolic traits, both in *Arabidopsis* and other species.

## Author contributions

STH, KB, TB, and RA conceived the study and wrote the paper. STH and KB performed the analyses. AL, EK, and TB contributed to statistical analyses and interpretations. RA supervised the study and aided interpretation of results. All authors read and approved the final manuscript.

## Acknowledgements

We are grateful to Dan Kliebenstein, Jinliang Yang, Jeffrey Ross-Ibarra, and anonymous reviewers for helpful comments and discussions that improved the manuscript. We thank Doug Speed for advice on the LDAK software and Marianne Slaten for assistance with HAPPI GWAS.

---

**Figure S1.**
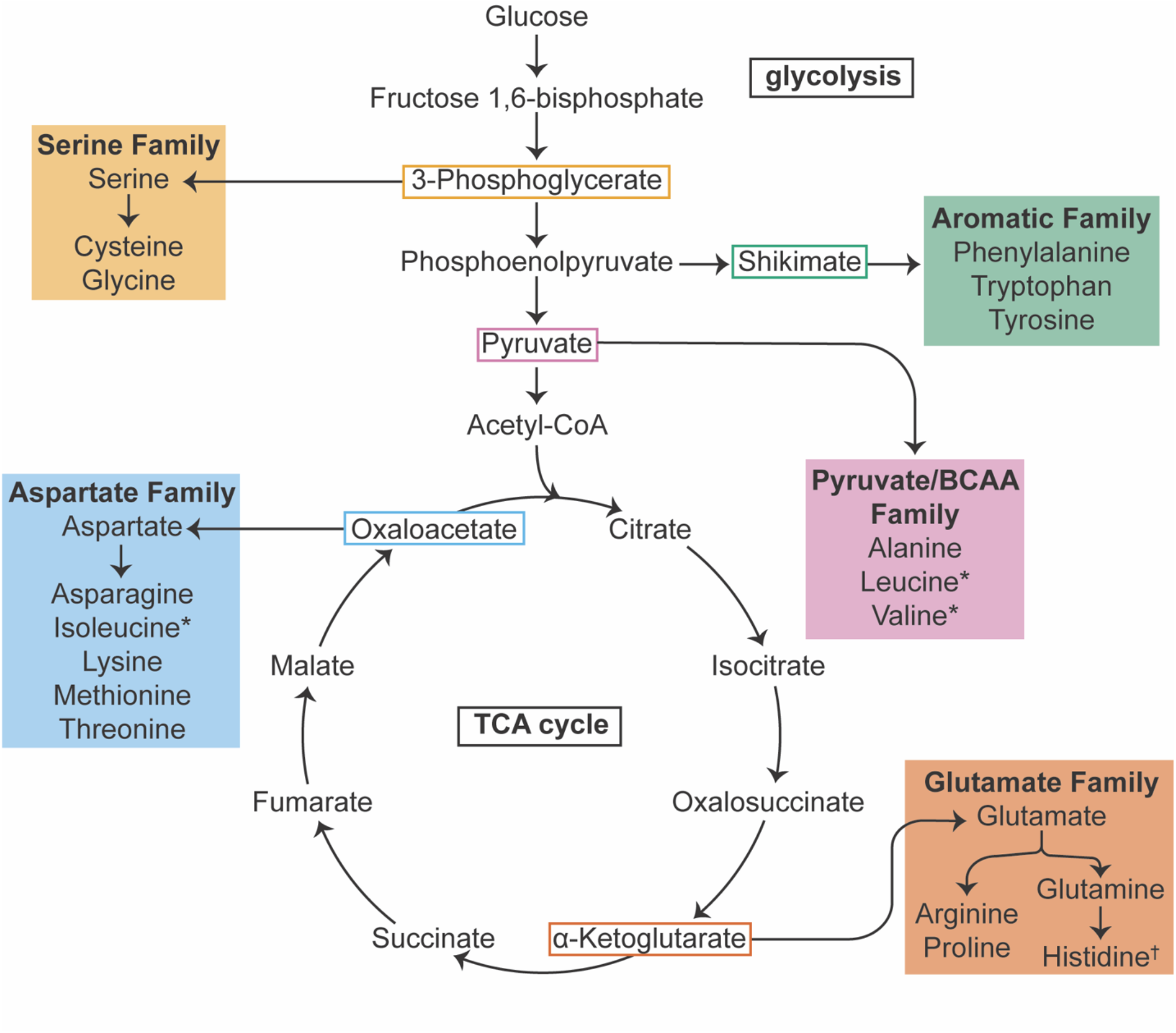
Biochemical relationships among amino acids. Colors indicate different amino acid families and boxes indicate the corresponding precursor. The branched-chain amino acids include Leu, Ile, Val are split across the Aspartate and Pyruvate family and therefore denoted with asterisks (*). Note that histidine (†) does not belong explicitly to the families identified here, but often is considered as part of the glutamate family.

**Figure S2.**
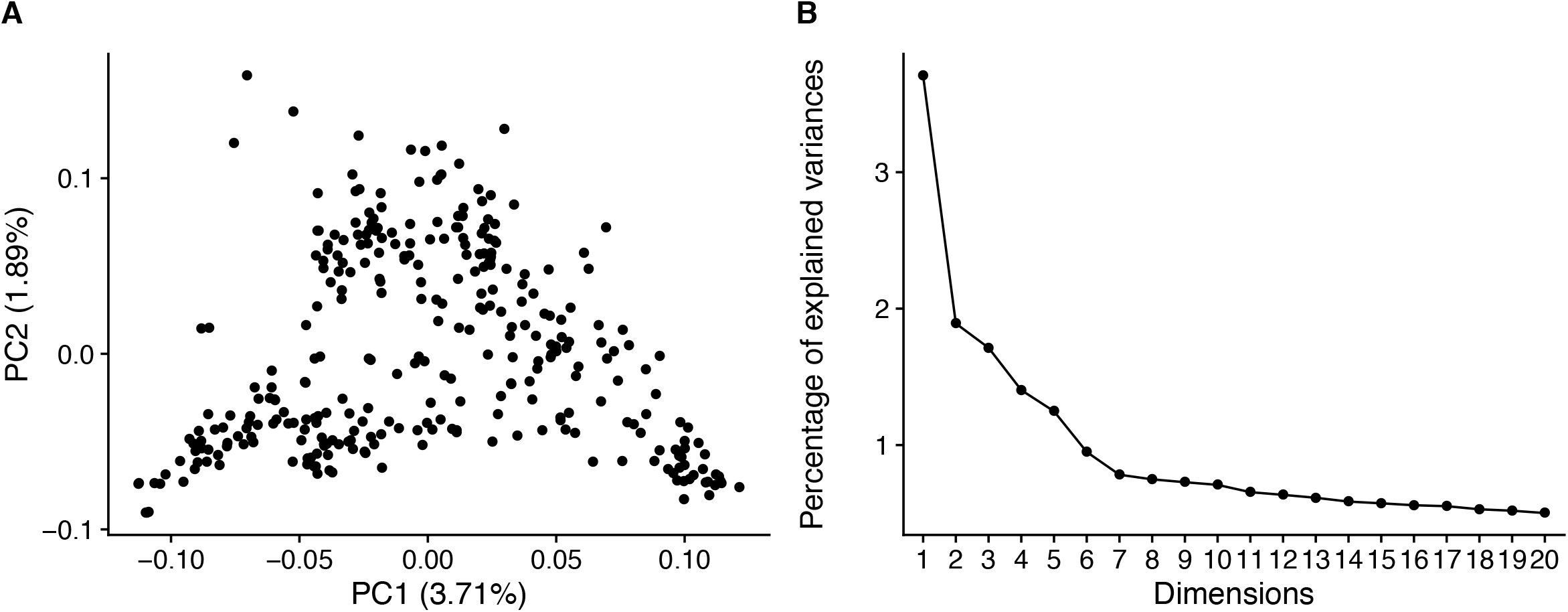
Principal component analysis (PCA) of genetic data for the 313 Arabidopsis accessions used in this study. (A) PCA scatterplot and percent variation explained for the first two principal components. (B) Screeplot showing the percent variance explained by each principal component.

**Figure S3.**
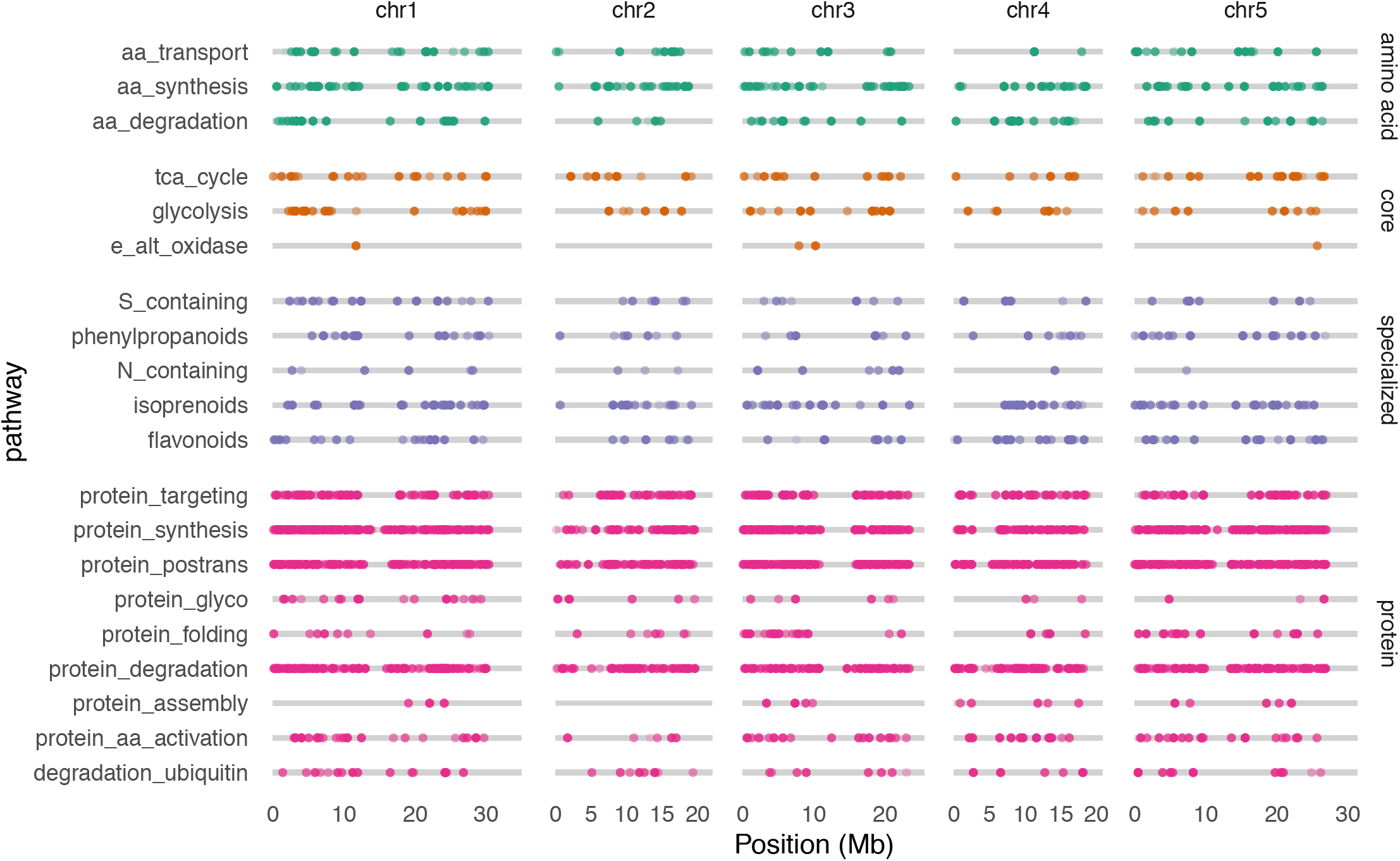
Genomic locations of pathway SNPs for each pathway. The x-axis indicates genomic position on each chromosome in megabases (Mb) and the y-axis indicates each pathway tested. Pathways are grouped into broad categories of metabolism (amino acid, core, specialized, and protein) and each point represents a SNP.

**Table S1.** List of seed free amino acid traits calculated from the quantification of 18 FAA and biochemical family affiliations. Strings of AA letter codes represent the sum of those AAs.

| Amino acids one letter code |  | Absolute levels |  | Relative levels to total |  | Biochemistry based metabolic ratios, grouped by AA families' affiliation |  |  |
| --- | --- | --- | --- | --- | --- | --- | --- | --- |
| Ala | A | AA-Abs<br>Total = Sum of 18 AA |  | AA/Total |  | Asp Family = Ile, Met, Thr, Asp Lys (IMTDK) |  | BCAA Family = Ile, Val, Leu (IVL)<br>Pyr Family=Leu, Ala, Val (LAV) |
| Asp | D | A |  | A/Total |  | D/IMTDK |  | I/IVL |
| Glu | E | D |  | D/Total |  | K/IMTDK |  | V/IVL |
| Phe | F | E |  | E/Total |  | M/IMTDK |  | L/IVL |
| Gly | G | F |  | F/Total |  | T/IMTDK |  | A/LAV |
| His | H | G |  | G/Total |  | IMTDK/D |  | L/LAV |
| Ile | I | H |  | H/Total |  |  |  | V/LAV |
| Lys | K | I |  | I/Total |  |  |  | I/IMTDK |
| Leu | L | K |  | K/Total |  | Glu Family = Glu, His, Pro, Arg, Gln (EHPRQ) |  | Shikimate (Aromatic) Fam = Trp, Phe,Tyr (WFY) |
| Met | M | L |  | L/Total |  | Q/EHPRQ |  | W/WFY |
| Pro | P | M |  | M/Total |  | E/EHPRQ |  | F/WFY |
| Gln | Q | P |  | P/Total |  | H/EHPRQ |  | Y/WFY |
| Arg | R | Q |  | Q/Total |  | P/EHPRQ |  |  |
| Ser | S | R |  | R/Total |  | R/EHPRQ |  | Ser Family = Ser, Gly (Cysteine-not detected -SG) |
| Thr | T | S |  | S/Total |  | EHPRQ/E |  | G/SG |
| Val | V | T |  | T/Total* |  |  |  | S/SG |
| Trp | W | V |  | V/Total |  |  |  |  |
| Tyr | Y | W |  | W/Total |  |  |  |  |
|  |  | Y |  | Y/Total |  |  |  |  |
|  |  | Total |  |  |  |  |  |  |
|  |  | IMTDK (Asp family) |  |  |  |  |  |  |
|  |  | IVL (BCAA family) |  |  |  |  |  |  |
|  |  | LAV (Pyr family) |  |  |  |  |  |  |
|  |  | EHPRQ (Glu family) |  |  |  |  |  |  |
|  |  | WFY (Shik family) |  |  |  |  |  |  |
|  |  | SG (Ser family) |  |  |  |  |  |  |

**Table S2.**
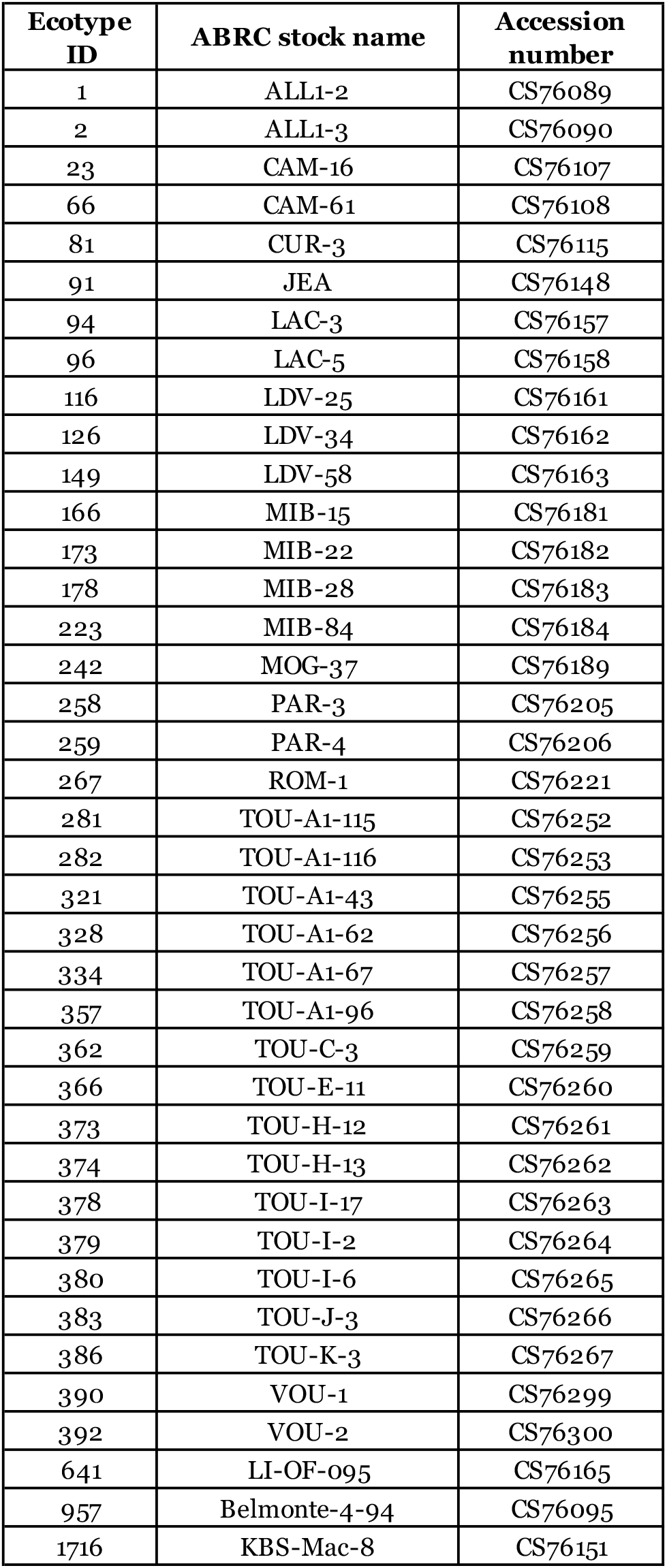
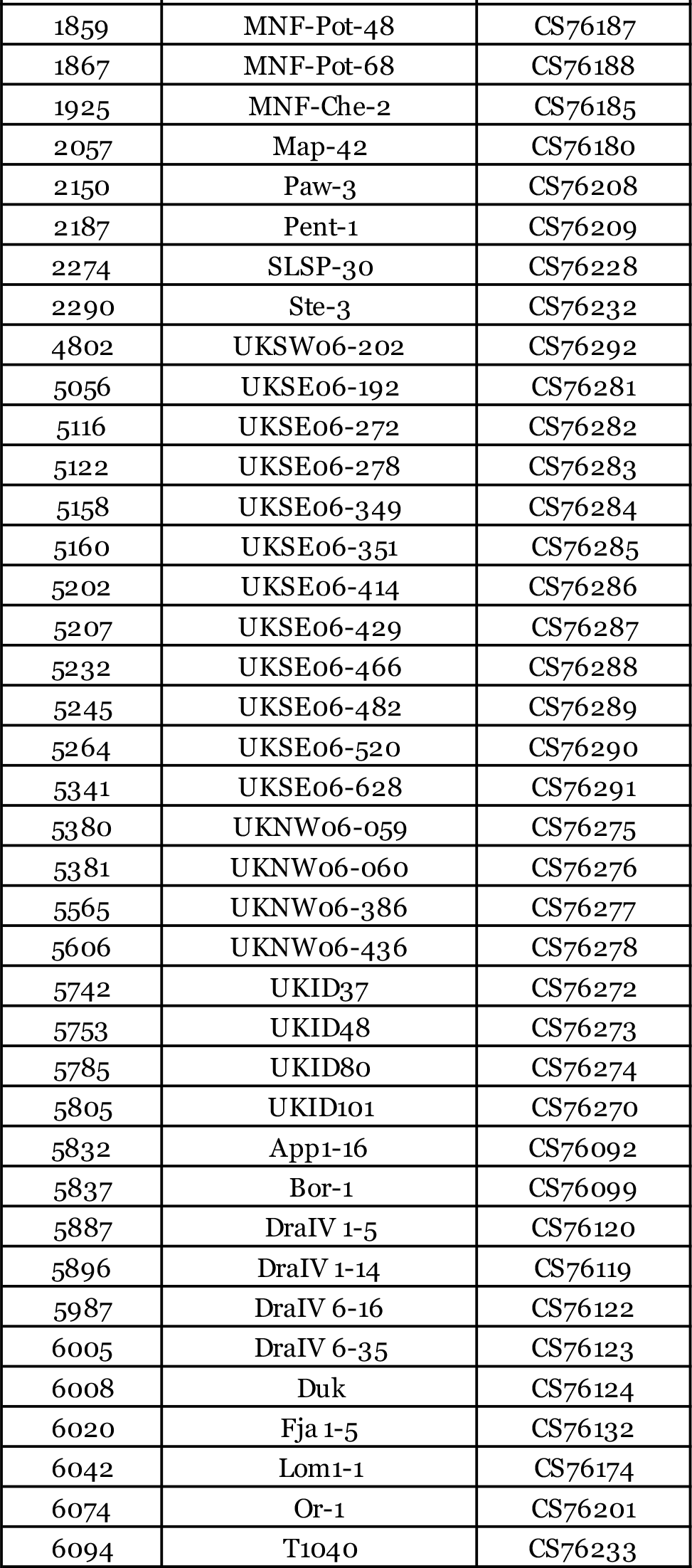
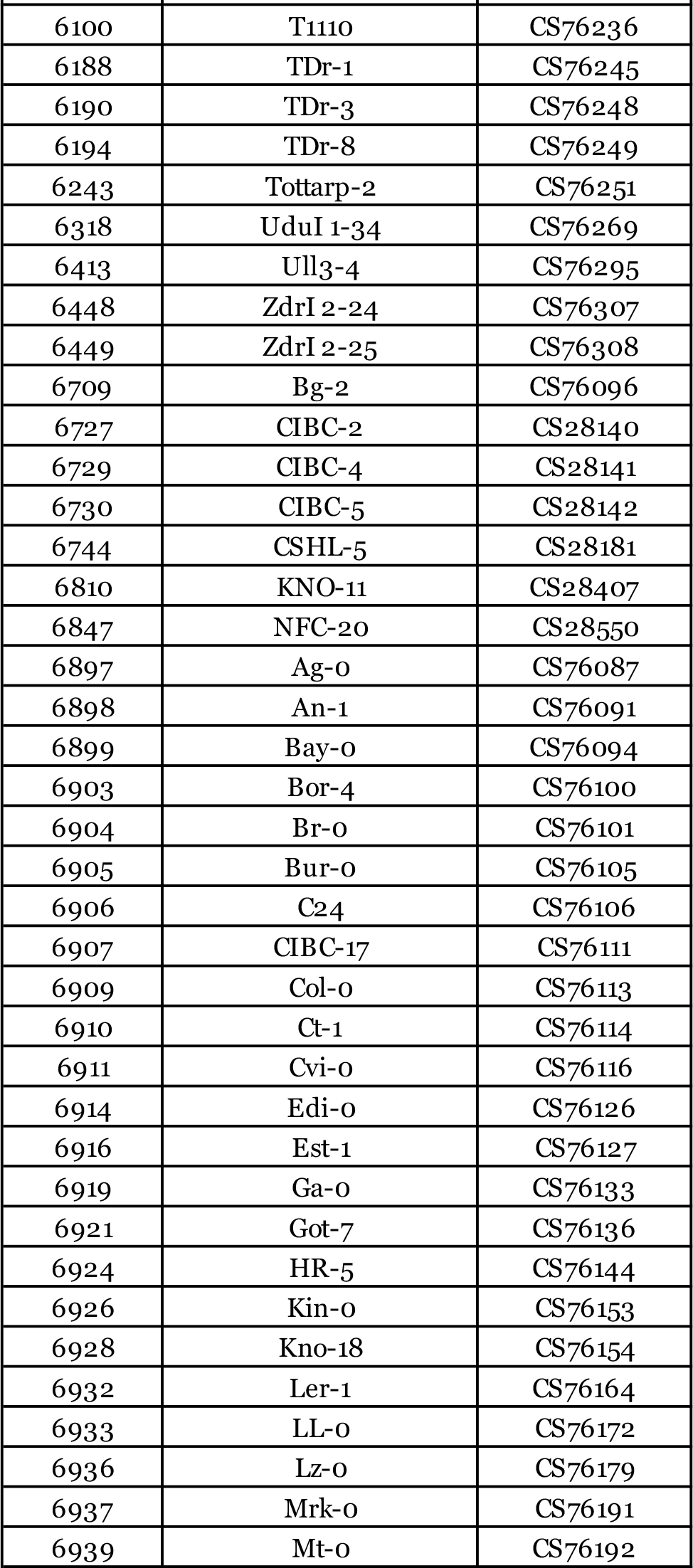
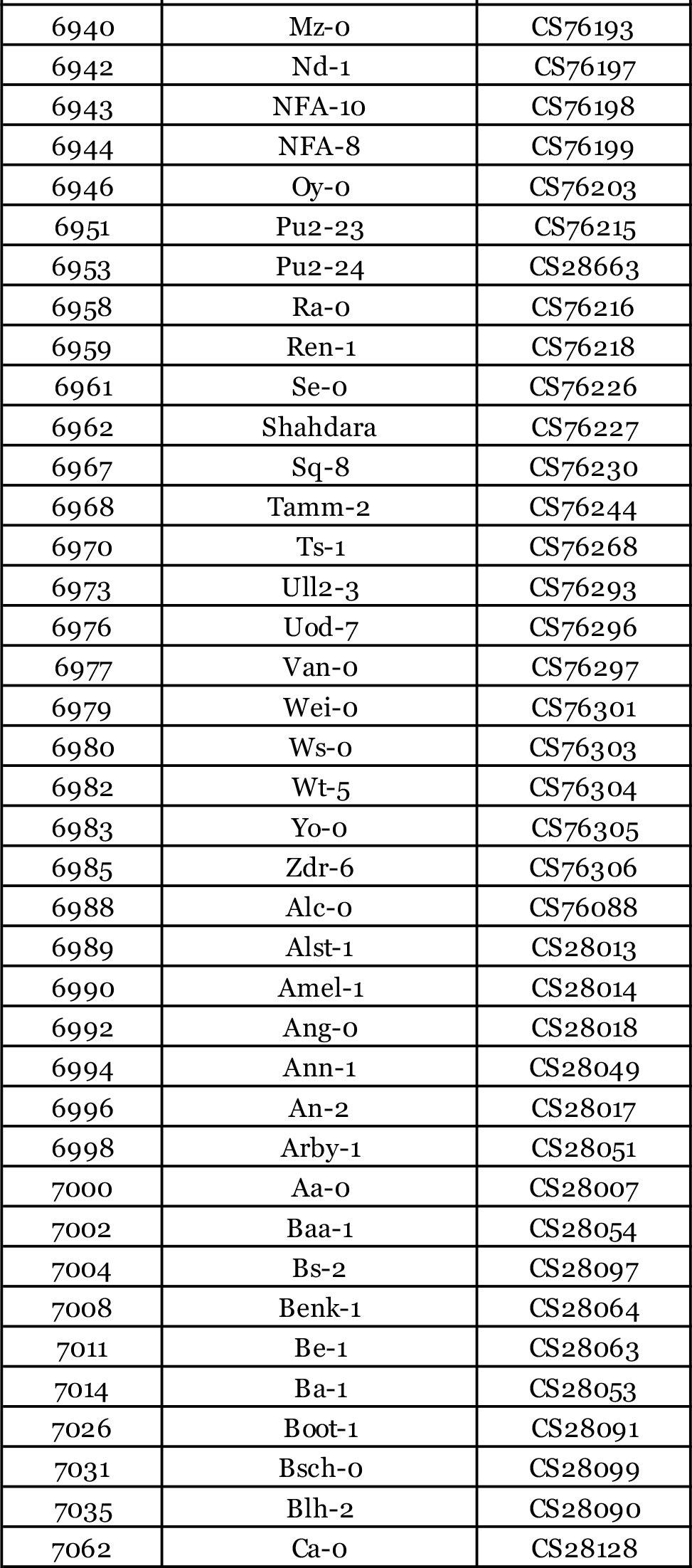
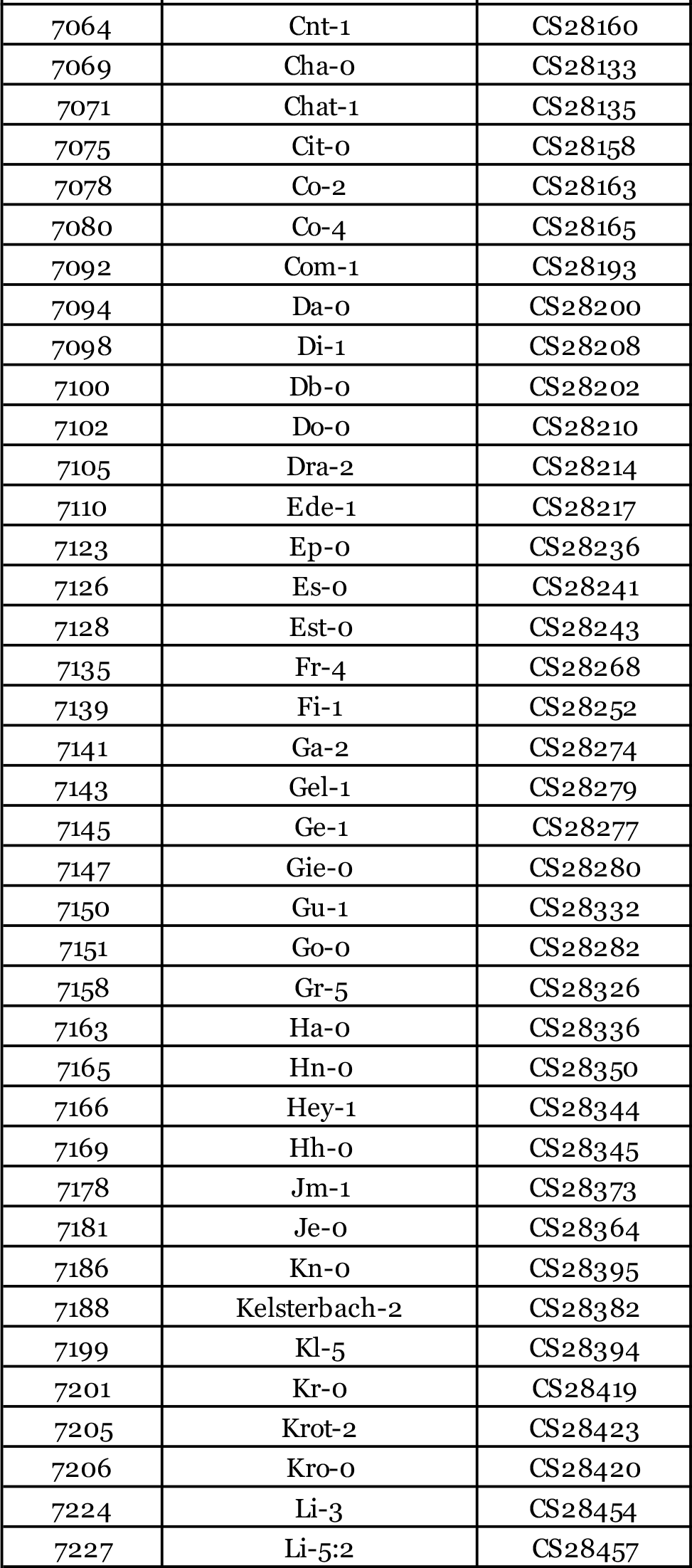
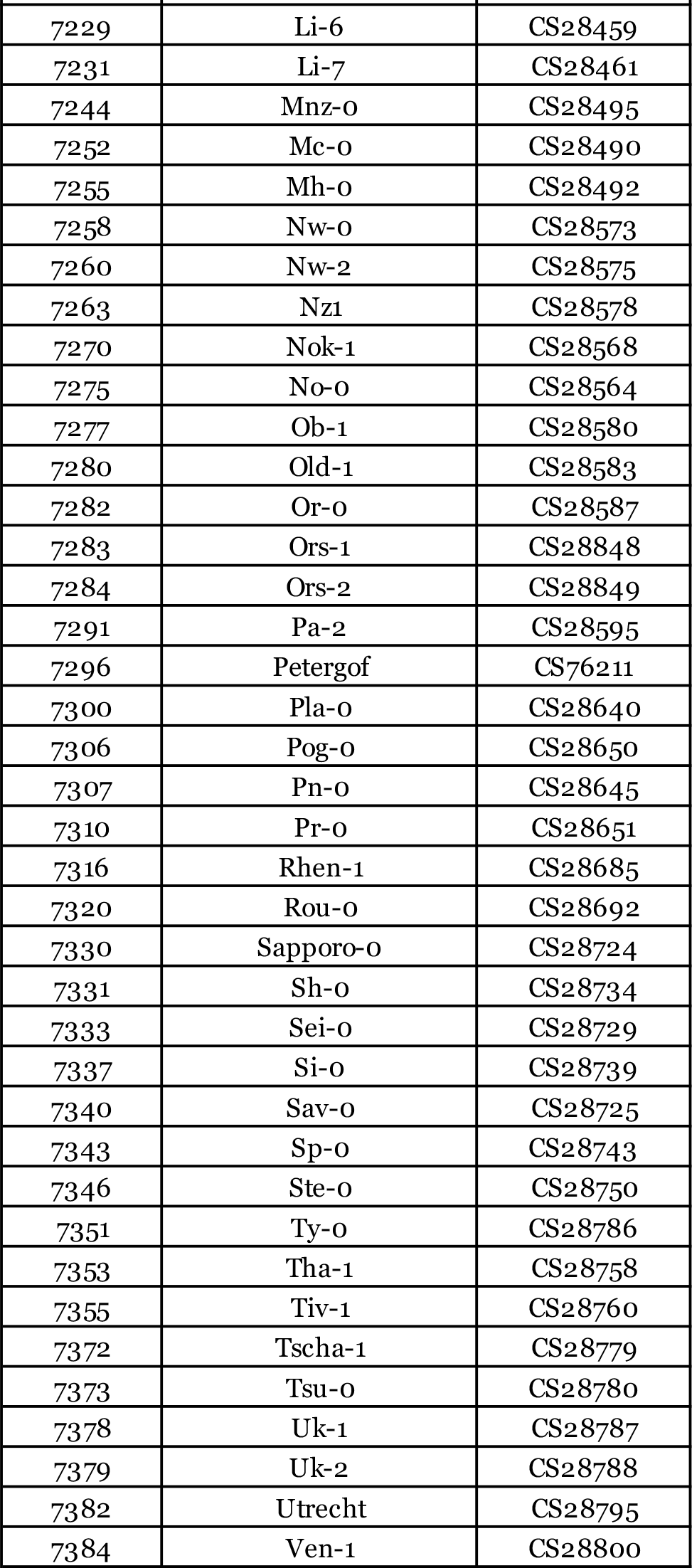
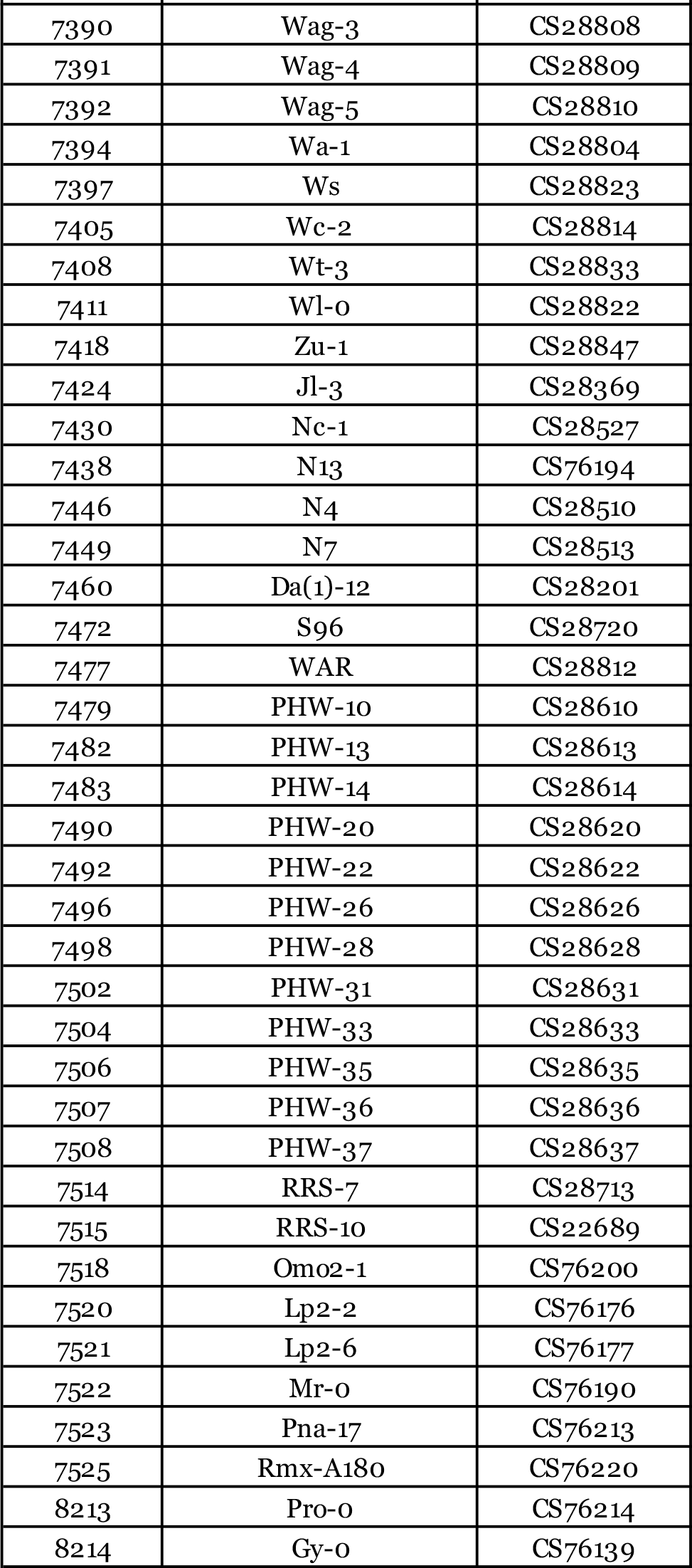
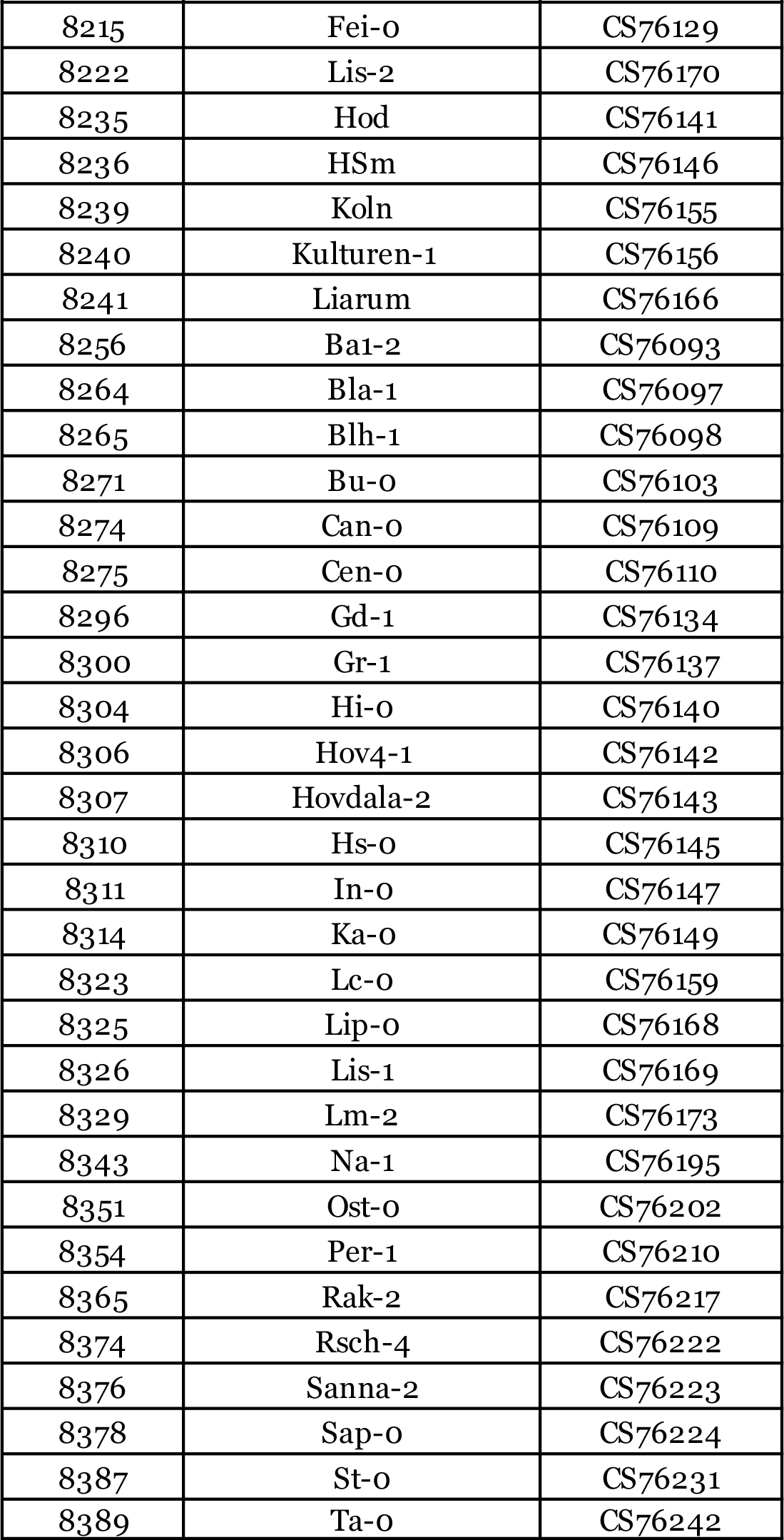
*Arabidopsis thaliana* accessions used in this study. (from Angelovici et al. Plant Cell 2013)

**Table S3.** Ranges for the number of genes and SNPs in 1000 null gene groups for each pathway. Null gene groups were sampled for each pathway by excluding SNPs within the pathway.

| **pathway** | **Number of genes** |  | **Number of SNPs** |  |
| --- | --- | --- | --- | --- |
|  | **minimum** | **maximum** | **minimum** | **maximum** |
| aa_degradation | 108 | 177 | 1080 | 1172 |
| aa_synthesis | 225 | 327 | 2068 | 2150 |
| aa_transport | 88 | 159 | 931 | 1017 |
| degradation_ubiquitin | 1924 | 2129 | 15866 | 16000 |
| e_alt_oxidase | 3 | 21 | 66 | 143 |
| flavonoids | 104 | 168 | 1054 | 1118 |
| glycolysis | 85 | 144 | 850 | 927 |
| isoprenoids | 189 | 269 | 1769 | 1859 |
| N_containing | 18 | 49 | 229 | 304 |
| phenylpropanoids | 84 | 139 | 839 | 909 |
| protein_aa_activation | 125 | 194 | 1223 | 1285 |
| protein_assembly | 25 | 65 | 310 | 392 |
| protein_degradation | 739 | 885 | 6363 | 6470 |
| protein_folding | 72 | 138 | 809 | 867 |
| protein_glyco | 40 | 81 | 452 | 529 |
| protein_postrans | 1030 | 1211 | 8725 | 8835 |
| protein_synthesis | 854 | 1023 | 7231 | 7342 |
| protein_targeting | 417 | 530 | 3670 | 3738 |
| S_containing | 70 | 119 | 725 | 800 |
| tca_cycle | 95 | 159 | 916 | 1003 |

